# Consensus Tree under the Ancestor-Descendant Distance is NP-hard

**DOI:** 10.1101/2023.07.17.549375

**Authors:** Yuanyuan Qi, Mohammed El-Kebir

**Author notes:** Authors’ address: Yuanyuan Qi,; Mohammed El-Kebir, University of Illinois Urbana-Champaign, Department of Computer Science, 201 N Goodwin Ave, Urbana, IL, USA, 61801.

## Abstract

Due to uncertainty in tumor phylogeny inference from sequencing data, many methods infer multiple, equally-plausible phylogenies for the same cancer. To summarize the solution space 𝒯 of tumor phylogenies, consensus tree methods seek a single best representative tree *S* under a specified pairwise tree distance function. One such distance function is the ancestor-descendant (AD) distance *d* (*T, T*′), which equals the symmetric difference of the transitive closures of the edge sets *E* (*T*) and *E* (*T*′). Here, we show that finding a consensus tree *S* for tumor phylogenies 𝒯 that minimizes the total AD distance ∑_*T* ∈𝒯_ *d* (*S, T*) is NP-hard.

## 1 INTRODUCTION

Cancer results from an evolutionary process during which somatic mutations accumulate in a population of cells [13]. To study tumor evolution, researchers apply phylogeny inference algorithms to sequencing data of tumors [15]. Due to uncertainty in tumor phylogeny inference from sequencing data, these methods typically yield multiple candidate trees 𝒯 for the same tumor [5, 14]. To summarize this solution space, several works have been proposed to infer a consensus tree *S* that best represents the set 𝒯 of candidate trees. More formally, these consensus tree methods typically employ a distance function *d* (*T*_1_, *T*_2_) that compares two trees *T*_1_ and *T*_2_, and seek a consensus tree *S* that minimizes the total distance ∑_*T* ∈𝒯_ *d* (*S, T*) [1, 4, 6–9, 11].

As many tumor phylogeny inference methods make the infinite-sites assumption (ISA), which states that each mutation is gained exactly once on the tree and never subsequently lost [12], consensus tree methods typically operate on mutation trees. These are rooted trees whose vertices are labeled by mutations that provide an equivalent representation of tumor phylogenies adhering to ISA (see Fig. 1a). For example, GraPhyC [7, 8] finds an optimal consensus tree based on the parent-child (PC) distance in polynomial time. Briefly, the PC distance of trees *T*_1_ and *T*_2_ is defined as the size of the symmetric difference between the edge sets *E* (*T*_1_) and *E* (*T*_2_). Aguse et al. [1] generalized the problem to identify multiple consensus trees under the PC distance, and Christensen et al. [2] considered a multiple-choice version of the problem to identify repeated evolutionary trajectories in cancer phylogeny cohort data. Although computationally tractable, Dinardo et al. [4] suggest the PC distance may not provide enough resolution for tumor phylogeny comparison.

**Fig. 1.**
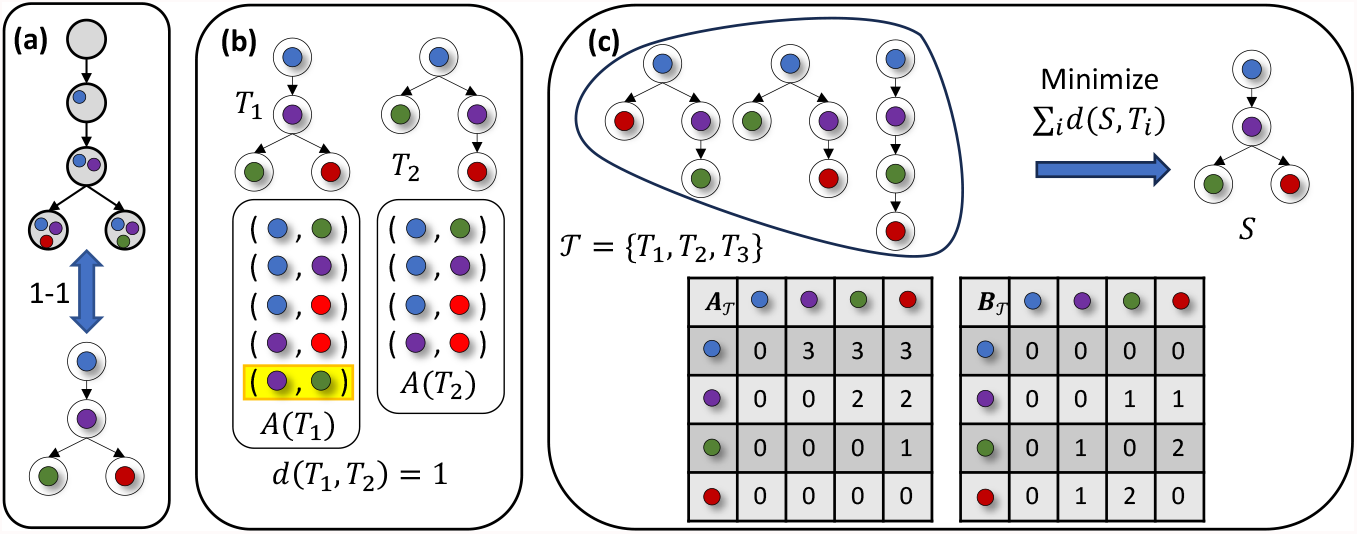
Overview of the Ancestor-Descendant Consensus Tree (ADCT) problem. (a) There is a bijection between phylogenies under the infinite sites assumption and mutation trees. (b) The ancestor-descendant distance *d* (*T*_1_, *T*_2_) of mutation trees *T*_1_ and *T*_2_ equals the size of the symmetric difference of the ancestor-descendant sets *A*(*T*_1_) and *A*(*T*_2_). Here, *d* (*T*_1_, *T*_2_) = 1 due to the unmatched pair of *A*(*T*_1_) highlighted in yellow. (c) In the ADCT problem, we seek a consensus mutation tree *S* that minimizes the sum of the distances to the input trees 𝒯.

Recently, Guang et al. [9] developed TuELiP, which uses integer linear programming to identify the optimal consensus tree with the ancestor-descendant (AD) distance, originally proposed in [7]. The AD distance equals the symmetric difference of the transitive closures of the edge sets *E* (*T*_1_) and *E* (*T*_2_), thus providing greater resolution to detect subtle differences between the two trees (see Fig. 1b). The hardness of the consensus tree problem under the AD distance is unknown (Fig. 1c).

In this paper, we show that finding the optimal consensus tree under the ancestor-descendant distance is NP-hard. Therefore, unlike the parent-child distance consensus tree problem, for which a polynomial-time algorithm exists, there is no polynomial-time algorithm for the consensus tree problem under the ancestor-descendant distance unless P=NP.

## 2 PROBLEM STATEMENT

We consider mutation trees *T*, which are rooted, vertex-labeled trees with vertex set *V* (*T*) and edge set *E* (*T*). Intuitively, each vertex *i* of a mutation tree *T* corresponds to a tumor clone comprised of the mutations that label the vertices of the unique path from *i* to the root of *T* (Fig. 1a). We write *i* ≺_*T*_ *j* if (i) vertex *i* is an ancestor of vertex *j* and (ii) *i* ≠ *j*. We write *i* ⊥_*T*_ *j* if vertices *i* and *j* occur on distinct root-to-leaf paths of *T*, i.e. *i* ⊀_*T*_ *j* and *j* ⊀_*T*_ *i*. While ⊥_*T*_ is symmetrical, i.e. *i* ⊥_*T*_ *j* if and only if *j* ⊥_*T*_ *i*, the relation ≺_*T*_ is not symmetrical. Neither ≺_*T*_ nor ⊥_*T*_ are reflexive, i.e. it does not hold that *i* ≺_*T*_ *i*, nor does it hold that *i* ⊥_*T*_ *i* for any vertex *i*. To compute the distance between two rooted trees *T*_1_ and *T*_2_ on the same vertex set, we compare the ancestor-descendant sets *A*(*T*_1_) and *A*(*T*_2_) of *T*_1_ and *T*_2_, respectively. More formally, *A*(*T*_1_) equals the transitive closure of *E* (*T*_1_).

### Definition 2.1.

The *ancestor-descendant (AD) set A*(*T*) of a rooted tree *T* consists of all ordered pairs (*i, j*) of vertices such that *i* is ancestor of *j*, i.e. *A*(*T*) = {(*i, j*) ∈ *V* (*T*) × *V* (*T*) | *i* ≺_*T*_ *j* }.

The ancestor-descendant (AD) distance *d* (*T*_1_, *T*_2_) equals the size of the symmetric difference of *A*(*T*_1_) and *A*(*T*_2_), more formally defined as follows. See Fig. 1b for an example.

### *Definition 2.2*.

Given two rooted trees *T*_1_, *T*_2_ on the same vertex set, the *ancestor-descendant (AD) distance d* (*T*_1_, *T*_2_) equals the size of the symmetric difference of *A*(*T*_1_) and *A*(*T*_2_), i.e. *d* (*T*_1_, *T*_2_) = |*A*(*T*_1_)\ *A*(*T*_2_) |+ |*A*(*T*_2_)\ *A*(*T*_1_)|.

This leads to the following problem.

### Problem 2.3

(Ancestor-Descendant Consensus Tree (ADCT)). *Given a multi-set* 𝒯 = {*T*_1_,…, *T*_*m*_} *of rooted trees on the same vertex set V*(𝒯), *find a rooted tree S on vertex set V*(𝒯) *such that the sum* 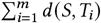 *of the distances from S to each input tree T* ∈ 𝒯 *is minimum*.

## 3 COMBINATORIAL CHARACTERIZATION

For any unordered pair {*i, j* } of distinct vertices in a mutation tree *T*, it must hold that *i* ≺_*T*_ *j, j* ≺_*T*_ *i* or *i* ⊥_*T*_ *j*. We indicate the first two cases using 1{*i* ≺_*T*_ *j* } such that 1{*i* ≺_*T*_ *j* } = 1 if *i* ≺_*T*_ *j* and 0 otherwise, and the third case using 1{*i* ⊥_*T*_ *j* } such that 1{*i* ⊥_*T*_ *j* } = 1 if *i* ⊥_*T*_ *j* and 0 otherwise. As such, the distance *d* (*T*_1_, *T*_2_) can be decomposed as follows.

### Lemma 3.1.

*The AD distance d* (*T*_1_, *T*_2_) *for trees T*_1_ *and T*_2_ *on the same vertex set* [*n*] *equals*

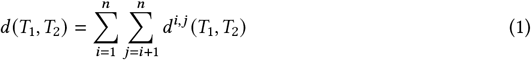

*where d*^*i,j*^ (*T*_1_, *T*_2_) *is the distance contributed by the unordered pair* {*i, j* } *of distinct vertices defined as*

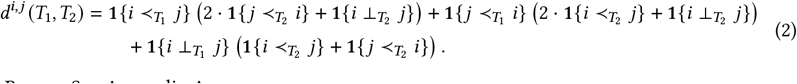

Proof. See Appendix A.

We can similarly decompose the total AD distance *d* (*S*, 𝒯) between a tree *S* and trees 𝒯 by first defining the ancestor-descendant matrix and the branching matrix as follows.

### *Definition 3.2*.

An *n* × *n* matrix *A*_𝒯_ = [*a*_*i,j*_] is an *ancestor-descendant matrix for trees* 𝒯 provided each entry *a*_*i,j*_ equals ∑_*T* ∈T_ 1{*i* ≺_*T*_ *j* }.

### *Definition 3.3*.

An *n* ×*n* matrix *B*_T_ = [*b*_*i,j*_] is an *branching matrix for trees* 𝒯 provided each entry *b*_*i,j*_ equals ∑_*T* ∈T_ 1{*i* ⊥_*T*_ *j* }.

While *A*_𝒯_ may not be symmetric, matrix *B*_𝒯_ is symmetric due to symmetry of the relation ⊥_*T*_. Moreover, the diagonal of both matrices consist of 0s. Finally, note that *a*_*i,j*_ + *a* _*j,i*_ + *b*_*i,j*_ = |𝒯| if *i* ≠ *j*. See Fig. 1c for an example.

### Lemma 3.4.

*The AD distance d* (*S*, 𝒯) *between a tree S and trees* 𝒯 *on the same vertex set* [*n*] *equals*

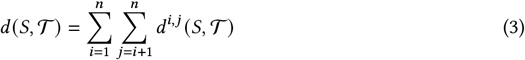

*where d*^*i,j*^ (*S*, 𝒯) *is the distance contributed by the unordered pair* {*i, j* } *of distinct vertices defined as*

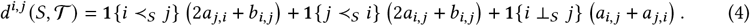

Proof. See Appendix A.

## 4 COMPLEXITY

Our main result is as follows.

### Theorem 4.1.

*The A**ncestor**-D**escendant* *C**onsensus* *T**ree* *problem is NP-hard*.

We show NP-hardness by giving a polynomial-time reduction from the MaxCliqe problem, defined as follows.

### Problem 4.1

(MaxCliqe). *Given an undirected graph G with n* = |*V* (*G*)| *vertices and m* = |*E* (*G*)| *edges, find a clique C* ⊆ *V* (*G*) *such that* |*C* | *is maximum*.

The MaxCliqe problem is NP-hard [3, 10]. In the following reduction, we assume the undirected graph *G* contains at least three vertices, i.e. *n* > 2. This assumption does not affect the hardness of the MaxCliqe problem. We impose an arbitrary ordering on the vertices *V* (*G*) such that *V* (*G*) = [*n*] = {1,…, *n*}. For each vertex *i* ∈ *V* (*G*), let *δ* (*i*) be the set of vertices adjacent to *i* in *G*, i.e. *δ* (*i*) = { *j* ∈ [*n*] | (*i, j*) ∈ *E* (*T*)}. Using the ordering, we further split the set *δ* (*i*) of neighboring vertices *δ* (*i*) into vertices *δ* ^*>*^ (*i*) = { *j* ∈ [*n*] | *j* ∈ *δ* (*i*), *i* < *j* } that are adjacent to *i* and occur after *i* in the ordering. The vertex set *V*(𝒯) of the corresponding ADCT problem instance includes 2*n* + 1 vertices, labeled {0, 1,…, *n, n* + 1,…, 2*n*}. Vertex 0 denotes a special vertex that is the shared root of all trees 𝒯, and a set {*n* + 1,…, 2*n*} of *n* vertices that forms a chain in all trees. We construct the following multi-set 𝒯 of rooted trees on the new vertex set *V*(𝒯) = {0,…, 2*n*}, with three types of trees — see Fig. 2b.

**Fig. 2.**
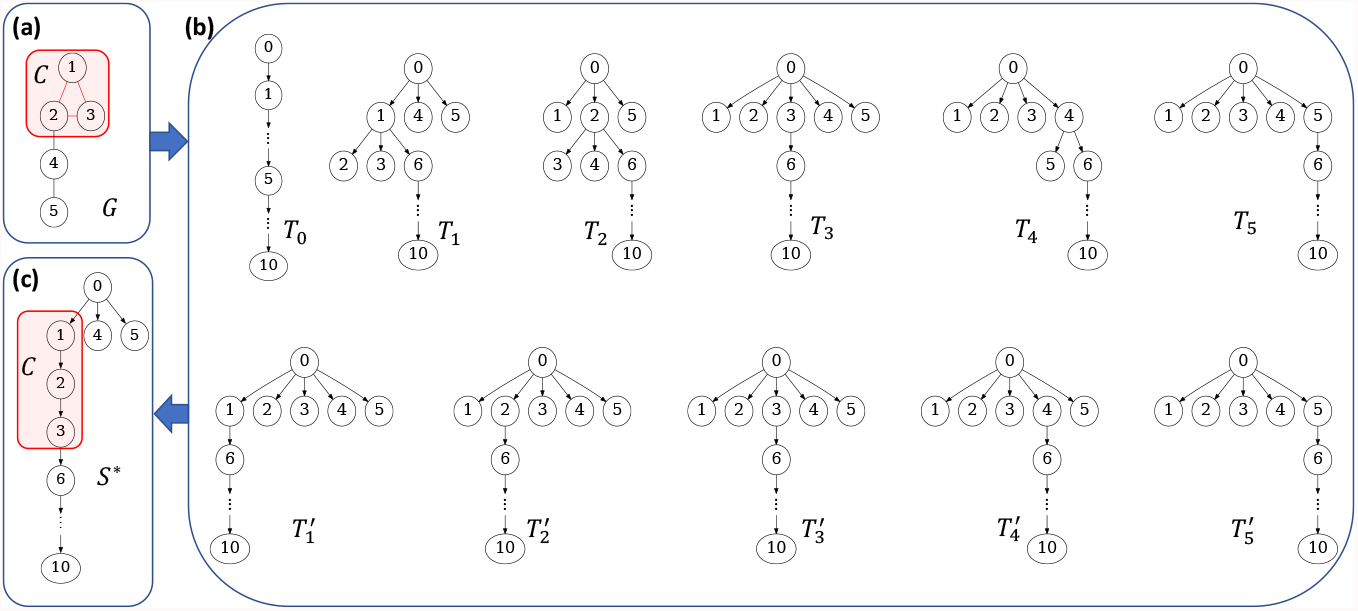
An example reduction from MaxClique to ADCT. (a) An undirected graph *G* with *n* = 5 vertices and *m* = 5 edges with a maximum clique *C* of size 3. Here, *δ* (2) = {1, 3, 4} and *δ* ^>^ (2) = {3, 4}. (b) The corresponding trees in 𝒯, with *n*^3^ − 2*n*^2^ + 2*n* − 3 = 82 copies of *T*_0_, *n*^2^ + 1 = 26 copies of *T*_*i*_ and one copy of 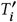 for each vertex *i* ∈ *V* (*G*). (c) The optimal consensus tree *S*\*. The vertices on the directed path between 0 and *n* + 1 = 6 indicate the maximum clique *C*.

First, let *T*_0_ be a chain tree whose vertices are in ascending order, i.e. *E* (*T*_0_) = {(*i, i* + 1) | 0 ≤ *i* < 2*n*}. The multi-set 𝒯_0_ is comprised of *n*^3^ − 2*n*^2^ + 2*n* − 3 copies of *T*_0_. Second, for each vertex *i* in the undirected graph *V* (*G*), let *T*_*i*_ be a tree rooted at 0. The edge set *E* (*T*_*i*_) consists of (i) edges from 0 to every vertex in *G* that is either at most *i* or is not adjacent to *i*, i.e. {(0, *j*) | *j* ∈ *V* (*G*)\ *δ* ^*>*^ (*i*)}; (ii) edges from 0 to every vertex is *G* that is greater than *i* and adjacent to *i*, i.e. {(*i, j*) | *j* ∈ *δ* ^*>*^ (*i*)}; (iv) the edge {(*i, n*+1)}; and (v) a chain from *n*+1 to 2*n* in ascending order, i.e. {(*j, j* +1) | *n* < *i* < 2*n*}. The multi-set 𝒯_*i*_ is comprised of *n*^2^ + 1 copies of *T*_*i*_. Third, for each vertex *i* ∈ *V* (*G*), let 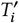 be a tree rooted at 0. The edge set 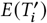 consists of (i) edges from 0 to every vertex in *G*, i.e. {(0, *j*) | *j* ∈ *V* (*G*)}; (ii) the edge {(*i, n*+1)}; and (iii) a chain from *n*+1 to 2*n* in ascending order, i.e. {(*j, j* +1) | *n*+1 ≤ *j* < 2*n*}. The multi-set 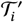 is comprised of only one copy of 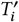.

The multi-set 𝒯 of trees corresponding to MaxCliqe instance *G* is comprised of the sum of multi-sets 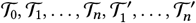. As such, the total number 𝒯 of trees equals 2*n*^3^ − 2*n*^2^ + 4*n* − 3. Clearly, the reduction can be completed in polynomial time. We have the following two lemmas characterizing the AD and branching matrix of 𝒯, respectively.

### Lemma 4.2.

*For any i, j* ∈ *V*(𝒯), *the entry a*_*i,j*_ *of the ancestor-descendant matrix A*_𝒯_ *equals:*

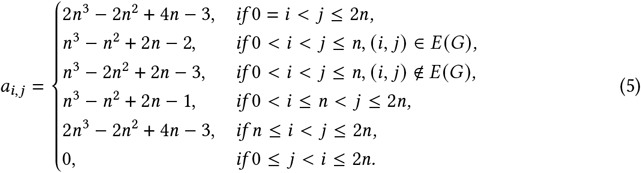

Proof. See Appendix A.

### Lemma 4.3.

*For any i, j* ∈ *V*(𝒯) *such that i* < *j, entries b*_*i,j*_ = *b* _*j,i*_ *of the branching matrix B*_𝒯_ *equal:*

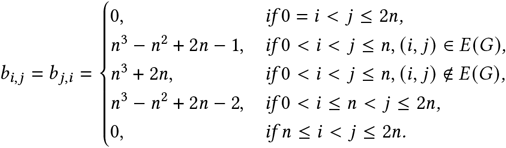

Proof. Recall that *a*_*i,j*_ + *a* _*j,i*_ + *b*_*i,j*_ = |𝒯|. Let *i, j* ∈ *V*(𝒯) such that *i* < *j*. Since *a* _*j,i*_ = 0 by Lemma 4.2, we have *a*_*i,j*_ + *b*_*i,j*_ = |𝒯|. As such, *b*_*i,j*_ = *b* _*j,i*_ = |𝒯| − *a*_*i,j*_. This lemma follows using the values of *a*_*i,j*_ from Lemma 4.2.

We prove the following lower bound on the distance *d* (*S*, 𝒯) of any tree *S* on vertex set *V*(𝒯).

### Lemma 4.4.

*If S isa tree on V*(𝒯) *then d* (*S*, 𝒯) *is at least* 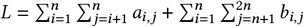.

Proof. Recall that for any pair (*i, j*) of vertices in any mutation tree *S*, exactly one of *i* ≺_*S*_ *j, j* ≺_*S*_ *i, i* ⊥_*S*_ *j* is true. Therefore, a trivial lower bound on *d*^*i,j*^ (*S*, T) can be obtained from Eq. (4): 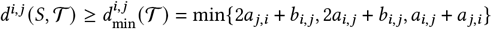. Note that *b*_*i,j*_ = *b* _*j,i*_ by Definition 3.3. As such, 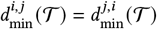. Using Lemma 4.2, if *i* < *j*, we have 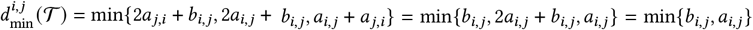. Furthermore, for *i* < *j*, by Lemma 4.3, we obtain

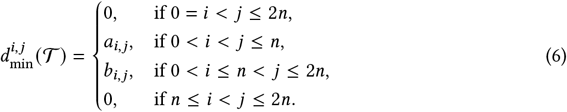

Plugging this into Eq. (3) of Lemma 3.4, we finally obtain

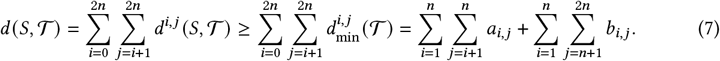

We define a *C*-constrained tree as follows — shown in Fig. 3a.

**Fig. 3.**
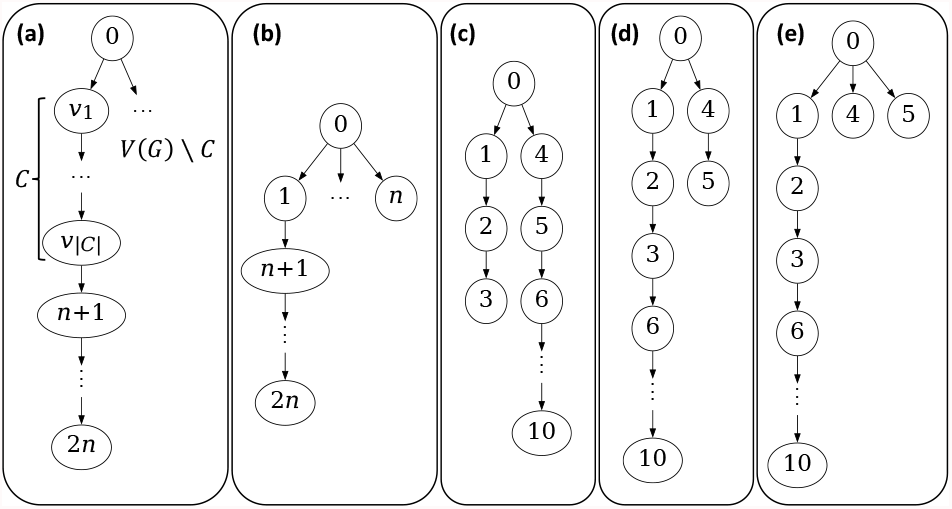
(a) The structure of a *C*-constrained tree *S*^*c*^ as well as an optimal consensus tree *S*\*. (b) The tree used to prove an upper bound on *d* (*S*\*, 𝒯) in Lemma 4.7. (c) An example tree based on the instance shown in Fig. 2 used in Lemma 4.12, where the chain {*n* + 1,…, 2*n*} is attached to vertex 5 which has smaller depth than vertex 3. (d) An example tree based on the instance shown in Fig. 2 used in Lemmas 4.12 and 4.14. In the former lemma, the chain {*n* + 1,…, 2*n*} is attached to vertex 5 which has higher depth than vertex 3. In the latter lemma, vertex 5 is a leaf whose parent is not 0. (e) An example tree based on the instance shown in Fig. 2 used in Lemma 4.14, where the parent of 5 is 0 instead of 4.

### *Definition 4.5*.

For vertices *C* = {*ν*_1_,…, *ν*_*k*_ } ⊆ *V* (*G*) of *G* such that *ν*_1_ < … < *ν*_*k*_, the *C-constrained tree S*_*C*_ has vertex set {0,…, 2*n*} such that (i) vertex 0 is the root, (ii) there is an edge (0, *i*) for each vertex *i* ∈ {1,…, *n*}\ *C*, and (iii) there is a chain 0 → *ν*_1_ → … → *ν*_*k*_ → *n* + 1 → … → 2*n*.

If *C* is a clique in *G* then the corresponding tree *S*_*C*_ induces the following distance *d* (*S*_*C*_, 𝒯).

### Lemma 4.6.

*For any clique C of size k of G, we have d* (*S*_*C*_, 𝒯) = *L* + *n*^2^ − *nk* + *k* (*k* − 1)/2.

Proof. Using Eq. (3) in Lemma 3.4, Lemmas 4.2, 4.3 and Eq. (6) in Lemma 4.4, we discuss the following six cases for the difference between *d*^*i,j*^ (*S*_*C*_, 𝒯) and 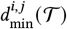 of vertices 0 ≤ *i* < *j* ≤ 2*n*.

First, we consider 0 = *i* < *j* ≤ 2*n*. Since *i* = 0 is the root vertex of *S*_*C*_. Therefore, it holds that 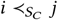. As such, 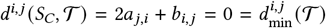. Second, we consider 0 < *i* < *j* ≤ *n* and *i, j* ∈ *C*. In this case, *i, j* are on the same branch in *S*_*C*_. Therefore, it holds that 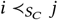. As such, *d*^*i,j*^ (*S*_*C*_, T) = 2*a* _*j,i*_ + *b*_*i,j*_ = *b*_*i,j*_. Since *C* is a clique, we have (*i, j*) ∈ *E* (*G*). Therefore, 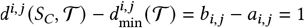. Third, we consider 0 < *i* < *j* ≤ *n*, and *i* ∉ *C* or *j* ∉ *C*. In this case, it holds that 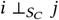. As such, 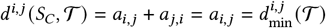. Fourth, we consider 0 < *i* ≤ *n* < *j* ≤ 2*n* and *i* ∉ *C*. In this case, it holds that 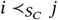. As such, 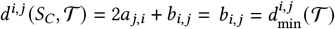. Fifth, we consider 0 < *i* ≤ *n* < *j* ≤ 2*n* and *i* ∉ *C*. In this case, it holds that 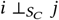. As such, *d*^*i,j*^ (*S*_*C*_, 𝒯) = *a*_*i,j*_ + *a* _*j,i*_ = *a*_*i,j*_. Therefore, 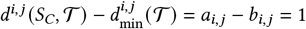. Sixth, we consider *n* ≤ *i* < *j* ≤ 2*n*. It holds that 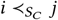. As such, 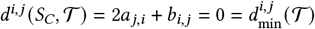.

Thus, only the second and fifth case have a non-zero value for 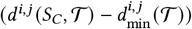. Putting everything together, we have that *d* (*S*_*C*_, 𝒯) − *L* equals

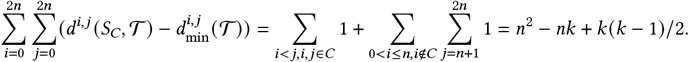

This proves the lemma.

Our goal now is to show that an optimal consensus tree *S*^*^ of the multi-set 𝒯 obtained from the undirected graph *G* is a *C*-constrained tree such that *C* is a clique of *G*. To this end, we establish the following useful upper bound on *d* (*S*^*^, 𝒯).

### Lemma 4.7.

*It holds that d* (*S*^*^, 𝒯) *is at most L* + *n*^2^ − *n*.

Proof. To prove the lemma we consider a tree *S* = (*V, E* (*S*)), *E* (*S*) = {(0, *i*) | 0 < *i* ≤ *n*}∪ {1, (*n* + 1)}∪ {(*i, i* + 1) | *n* < *i* < 2*n*} as shown in Fig. 3b. By Lemmas 3.4, 4.2 and 4.3, we have 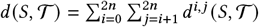 where

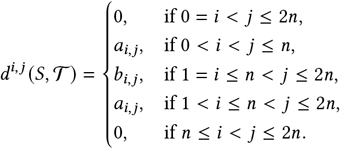

Observe that *a*_*i,j*_ = *b*_*i,j*_ + 1 for 0 < *i* ≤ *n* < *j* ≤ 2*n* in Lemmas 4.2 and 4.3. Using the lower bound *L* established in Lemma 4.4, we have

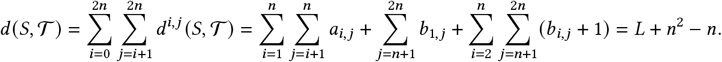

Hence, *d* (*S*^*^, 𝒯) ≤ *d* (*S*, 𝒯) = *L* + *n*^2^ − *n*.

We now reason about the topology of *S*^*^. The following lemma shows that *j* cannot be an ancestor of *i* in *S*^*^ if *i* < *j*.

### Lemma 4.8.

*For any pair* (*i, j*) *of vertices such that* 0 ≤ *i* < *j* ≤ 2*n, either i* ≺_*S*_* *j or i* ⊥_*S*_* *j*.

Proof. See Appendix A.

Our reduction enforces that 0 is the root of *S*^*^ and that 2*n* is a leaf.

### Lemma 4.9.

*Vertex* 0 *is the root of S*^*^.

Proof. Suppose for a contradiction that 0 < *j* ≤ 2*n* is the root of *S*^*^. Consider vertex 0. Since *j* is the root, it holds that *j* ≺_*S*_* 0. However, since 0 < *j*, by Lemma 4.8, it must hold that either 0 ≺_*S*_* *j* or 0 ⊥_*S*_* *j*, yielding a contradiction. Hence, vertex 0 must be the root of *S*^*^.

### Lemma 4.10.

*The subgraph of S*^*^ *induced by vertices* {0,…, *n*} *forms a tree*.

Proof. It suffices to prove that no vertex {*n* + 1,…, 2*n*} is an ancestor of a vertex {1,…, *n*} in *S*^*^. Suppose for a contradiction there exist vertices 0 < *i* ≤ *n* < *j* ≤ 2*n* such that *j* ≺_*S*_* *i*. Since *i* < *j*, by Lemma 4.8, it must hold that either *i* ≺_*S*_* *j* or *i* ⊥_*S*_* *j*, yielding a contradiction.

Moreover, vertices {*n* + 1,…, 2*n*} form a chain from *n* + 1 to 2*n* in ascending order as shown by the following lemma.

### Lemma 4.11.

*For any pair* (*i, j*) *of vertices such that n* < *i* < *j* ≤ 2*n, it holds that i* ≺_*S*_* *j*.

Proof. Suppose for a contradiction that *u* ⊀_*S*_* *v* for some *n* < *u* < *v* ≤ 2*n*. By Lemma 4.8, we have *v* ⊀_*S*_* *u*. Therefore, *u* ⊥_*S*_* *v*, i.e. *u* and *v* are branched in *S*^*^. By Eq. (3) in Lemma 3.4 and Lemma 4.2, we have *d*^*u,v*^ (*S*^*^, T) = *a*_*u,v*_. As such,

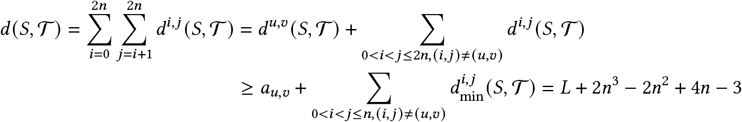

Since 2*n*^3^ − 2*n*^2^ + 4*n* − 3 > *n*^2^ − *n*, Lemma 4.7, which states that *d* (*S*^*^, 𝒯) ≤ *L* + *n*^2^ − *n*, implies *S*^*^ is not an optimal consensus tree, a contradiction.

The root vertex *r* of a mutation tree *S* has depth_*S*_ (*r*) = 0, and every vertex *v* ≠ *r* with parent *u* has depth_*S*_ (*v*) = depth_*S*_ (*u*)+ 1. In Lemma 4.11, we have shown that the chain {*n* + 1,…, 2*n*} remains intact in an optimal consensus tree *S*^*^. In the following lemma, we will show that this chain is attached to a maximum-depth vertex among {0,…, *n*} in *S*^*^.

### Lemma 4.12.

*The parent of vertex n* + 1 *in S*^*^ *is a vertex in the set* {0,…, *n*} *with maximum depth*.

Proof. Let *i* be the parent of vertex *n* + 1. Lemma 4.11 states that the chain *n* + 1 → … → 2*n* is kept intact in *S*^*^. This means that *i* must be in {0,…, *n*}. Suppose for a contradiction that depth_*S*_* (*i*) does not have the maximum depth among vertices {0,…, *n*}. Therefore, there is a vertex 0 ≤ *j* ≤ *n* such that depth_*S*\*_ (*j*) > depth_*S*\*_ (*i*) — as illustrated in Fig. 3c. Let *P*_*i*_ be the unique path from 0 to *i*. Let *P*_*j*_ be the unique path from 0 to *j*. Since depth_*S*\*_ (*j*) > depth_*S*\*_ (*i*), we have |*V* (*P*_*i*_)| < |*V* (*P*_*j*_)|. We remove the chain and re-attach it to the higher-depth vertex *j*, yielding *S* = (*V, E* (*S*)), where *E* (*S*) = (*E* (*S*^*^)\ {(*i, n* + 1)}) ∪ {(*j, n* + 1)} as shown in Fig. 3d. By Lemma 4.10, *S* is a tree. We will show that *d* (*S*, 𝒯) < *d* (*S*^*^, 𝒯) by distinguishing four cases regarding the placement of vertices 0 ≤ *u* < *v* ≤ 2*n*.

First, we consider 0 ≤ *u* < *v* ≤ *n* or *n* < *u* < *v* ≤ 2*n*. In the former case, *u, v* are located outside the chain and in the latter case, inside the chain. In both cases, the relation between *u* and *v* is the same in both *S*^*^ and *S*. As such, *d*^*u,v*^ (*S*^*^, 𝒯) − *d*^*u,v*^ (*S*, 𝒯) = 0 by Eq. (3) in Lemma 3.4. Second, we consider 0 ≤ *u* ≤ *n* < *v* ≤ 2*n* and *u* ∈ *V* (*P*_*i*_) ∩ *V* (*P*_*j*_). The relation between *u* and *v* also stays the same, and *u* is an ancestor of *v* in both *S*^*^ and *S*. Similar to the previous case, we have *d*^*u,v*^ (*S*^*^, 𝒯) − *d*^*u,v*^ (*S*, 𝒯) = 0. Third, we consider 0 ≤ *u* ≤ *n* < *v* ≤ 2*n* and *u* ∈ *V* (*P*_*i*_) \ *V* (*P*_*j*_). Thus, *u* is an ancestor of *v* in *S*^*^, however, they are branched in *S*. By Eq. (3) in Lemma 3.4, Lemma 4.2 and Lemma 4.3, *d*^*u,v*^ (*S*^*^, *T*) − *d*^*u,v*^ (*S, T*) = *b*_*u,v*_ − *a*_*u,v*_ = −1. Fourth, we consider 0 ≤ *u* ≤ *n* < *v* ≤ 2*n* and *u* ∈ *V* (*P*_*j*_) \ *V* (*P*_*i*_). Thus, *u* is an ancestor of *v* in *S*, however, they are branched in *S*^*^. Similarly to the previous case, we have *d*^*u,v*^ (*S*^*^, 𝒯) − *d*^*u,v*^ (*S*, 𝒯) = *a*_*u,v*_ − *b*_*u,v*_ = 1. Therefore,

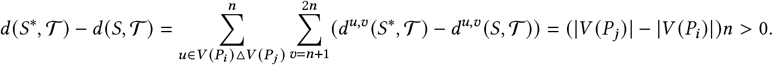

Note that Δ, indicates the symmetric difference. This contradicts that *S*^*^ is optimal.

We have that non-adjacent vertices *i, j* in *G* must be branched in an optimal consensus tree *S*^*^.

### Lemma 4.13.

*For any pair i, j* ∈ *V* (*G*) *of distinct vertices where* (*i, j*) ∉ *E* (*G*), *it holds that i* ⊥_*S*_* *j*.

Proof. Suppose for a contradiction there exist vertices *u, v* ∈ *V* (*G*) such that (*u, v*) ∉ *E* (*G*) and *u* /⊥_*S*_* *v*. WLOG, we assume *u* < *v*. By Lemma 4.8, either *u* ≺_*S*_* *v* or *u* ⊥_*S*_* *v*. Therefore, it holds that *u* ≺_*S*_* *v*. By Eq. (3) in Lemma 3.4 and Lemma 4.2, *d*^*u,v*^ (*S, T*) = *b*_*u,v*_. As such, *d* (*T*, 𝒯) equals

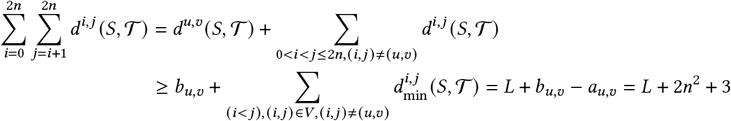

Since 2*n*^2^ + 3 > *n*^2^ − *n*, Lemma 4.7, which states that *d* (*S*^*^,𝒯) ≤ *L* + *n*^2^ − *n*, implies *S*^*^ is not an optimal consensus tree, a contradiction.

In the following lemma we show that *S*^*^ is a star tree except for one linear branch containing a subset *C* ⊆ {1,…, *n*} of vertices and terminating with the chain {*n* + 1,…, 2*n*} (Fig. 3a,e).

### Lemma 4.14.

*Let C be the vertices on the unique path from vertex* 0 *to vertex n* + 1, *excluding* 0 *and n* + 1. *Then, vertex* 0 *is the parent of all vertices* {1,…, *n*}\ *C in S*^*^.

Proof. Suppose for a contradiction that there are vertices in {1,…, *n*}\ *C* whose parents are not 0 in *S*^*^. Among these vertices, consider a leaf vertex *i*. Let vertex *j* ≠ 0 be the parent of *i* (see Fig. 3d where *i* = 5 and *j* = 4). Recall that by Lemma 4.9, vertex 0 is the root of *S*^*^. Let *V*_*i*_ be the vertices on the unique path from vertex 0 to *i*, excluding 0 and *i*. By Lemma 4.8, *V*_*i*_ consists of vertices *u* such that *u* < *i*. Consider the tree *S* where we attach vertex *i* to the root 0, i.e. *E* (*S*) = *E* (*S*^*^)\ {(*j, i*)} ∪ {(0, *i*)}. See Fig. 3e for an example. To compute *d*^*u,v*^ (*S*^*^, 𝒯) − *d*^*u,v*^ (*S*, 𝒯), we distinguish two cases for vertices 0 ≤ *u* < *v* ≤ 2*n*. First, we consider *v* = *i* and *u* ∈ *V*_*i*_. Thus, *u* ≺_*S*_* *v*. By the contrapositive of Lemma 4.13, we have (*u, i*) ∈ *E* (*G*) for any *u* ∈ *V*_*i*_. Since *u* ⊥_*S*_ *v*, by Eq. (3) in Lemma 3.4, Lemma 4.2 and Lemma 4.3, we have *d*^*u,v*^ (*S*^*^, 𝒯) − *d*^*u,v*^ (*S*, 𝒯) = *b*_*u,v*_ − *a*_*u,v*_ = 1. Second, we consider the case where *v* ≠ *i* or *u* ∉ *V*_*i*_. The relationship between *u* and *v* is the same in *S* and *S*^*^. As such, *d*^*u,v*^ (*S*^*^, 𝒯) − *d*^*u,v*^ (*S*, 𝒯) = 0 by Eq. (3) in Lemma 3.4. Therefore,

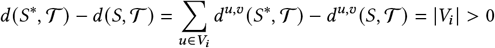

This contradicts that *S*^*^ is optimal and thus proves the lemma.

Finally, we show that vertices *C* of *S*^*^ is indeed a clique of *G*.

### Lemma 4.15.

*The vertices C of S*^*^ *on the unique path from vertex* 0 *to vertex n* + 1, *excluding* 0 *and n* + 1, *form a clique in G*.

Proof. By Lemma 4.9, vertex 0 is the root of *S*^*^. Therefore, for any *i, j* ∈ *C*, we have *i* /⊥_*S*_* *j*. By the contrapostive of Lemma 4.13, (*i, j*) ∈ *E* (*G*) for all *i, j* ∈ *C*. Hence, *C* is a clique of *G*.

### Corollary 4.16.

*Any optimal consensus tree S*^*^ *is a C-constrained tree such that C is a clique of G*.

### Lemma 4.17.

*For any subset C* ⊆ *V* (*G*) *of vertices, the C-constrained tree S*_*C*_ *is an optimal consensus tree of* T *if and only if C is a maximum clique in G*.

Proof. (⇒) Let *S*_*C*_ be an optimal *C*-constrained consensus tree. By Lemma 4.15, we know that *C* is a clique. Let |*C* | = *k*. By Lemma 4.6, we have *d* (*S*_*C*_, 𝒯) = *L* + *n*^2^ − *nk* + *k* (*k* − 1)/2. Suppose for a contradiction that *C* is not a maximum clique of *G*. By our premise, there must exist another clique *C*′ such that |*C*′ | = *ℓ* > *k* = |*C* |. Let 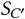 be the corresponding *C*′-constrained tree following Definition 4.5. By Lemma 4.6, we have *d* (*S*_*C*_′, 𝒯) = *L* + *n*^2^ − *nℓ* + *ℓ* (*ℓ* − 1)/2. Since *n* ≥ *ℓ* ≥ *k* + 1, 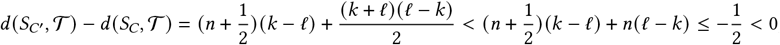 which contradicts that *S*_*C*_ is optimal.

(⇐) Let *C* be a maximum clique of *G* such that |*C* | = *k*. Suppose for a contradiction that the corresponding *C*-constrained tree *S*_*C*_ is not an optimal consensus tree of 𝒯. By Lemma 4.6, we have *d* (*S*_*C*_, 𝒯) = *L* + *n*^2^ − *nk* + *k* (*k* − 1)/2. Therefore, by Corollary 4.16, there exists an optimal *C*′-constrained consensus tree *S*_*C*_′, where |*C*′ | = *ℓ*, such that the distance *d* (*S*_*C*′_, 𝒯) is strictly less than *d* (*S*_*C*′_, 𝒯). By Lemma 4.6, *d* (*S*_*C*′_, 𝒯) = *L* + *n*^2^ − *nℓ* + *ℓ* (*ℓ* − 1)/2. We have

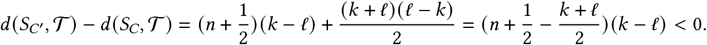

Since *k, ℓ* ≤ *n*, we have (*k* + *ℓ*)/2 ≤ *n*. This implies that *k* − *ℓ* < 0, which contradicts that *C* is a maximum clique of size *k*.

## 5 DISCUSSION

In this work, we demonstrated the NP-hardness of the consensus tree problem under the ancestor-descendant (AD) distance. While the problem of finding a maximum clique for a graph with *n* vertices is hard to approximate within a factor of *O* (*n*^1−*∈*^) for any real number ∈ > 0 unless P=NP [16], our reduction is not approximation-factor preserving. As such, one might be able to achieve better approximation factors for the consensus tree problem under the AD distance, including possibly constant factors. We will investigate this in future work.

## Supporting information

Supplementary Proofs

