## Supplementary Proofs for "Consensus Tree under the Ancestor-Descendant Distance is NP-hard"

### A SUPPLEMENTARY PROOFS

(Main Text) Lemma 3.1. *The AD distance $d(T_1, T_2)$ for trees $T_1$ and $T_2$ on the same vertex set $[n]$ equals*

$$d(T_1, T_2) = \sum_{i=1}^{n} \sum_{j=i+1}^{n} d^{i,j}(T_1, T_2) \tag{8}$$

*where $d^{i,j}(T_1, T_2)$ is the distance contributed by the unordered pair $\{i, j\}$ of distinct vertices defined as*

$$\begin{aligned} d^{i,j}(T_1, T_2) = {} & \mathbf{1}\{i \prec_{T_1} j\} \left(2 \cdot \mathbf{1}\{j \prec_{T_2} i\} + \mathbf{1}\{i \perp_{T_2} j\}\right) + \mathbf{1}\{j \prec_{T_1} i\} \left(2 \cdot \mathbf{1}\{i \prec_{T_2} j\} + \mathbf{1}\{i \perp_{T_2} j\}\right) \\ & + \mathbf{1}\{i \perp_{T_1} j\} \left(\mathbf{1}\{i \prec_{T_2} j\} + \mathbf{1}\{j \prec_{T_2} i\}\right). \end{aligned} \tag{9}$$

Proof. By Definition 2.2, we have

$$\begin{aligned} d(T_1, T_2) &= |A(T_1) \setminus A(T_2)| + |A(T_2) \setminus A(T_1)| \\ &= \sum_{i=1}^{n} \sum_{j=1}^{n} \mathbf{1}\{i \prec_{T_1} j\}\mathbf{1}\{i \not\prec_{T_2} j\} + \sum_{i=1}^{n} \sum_{j=1}^{n} \mathbf{1}\{i \prec_{T_2} j\}\mathbf{1}\{i \not\prec_{T_1} j\} \\ &= \sum_{i=1}^{n} \sum_{j=i+1}^{n} \left(\mathbf{1}\{i \prec_{T_1} j\}\mathbf{1}\{i \not\prec_{T_2} j\} + \mathbf{1}\{j \prec_{T_1} i\}\mathbf{1}\{j \not\prec_{T_2} i\}\right) \\ &\quad + \sum_{i=1}^{n} \sum_{j=i+1}^{n} \left(\mathbf{1}\{i \prec_{T_2} j\}\mathbf{1}\{i \not\prec_{T_1} j\} + \mathbf{1}\{j \prec_{T_2} i\}\mathbf{1}\{j \not\prec_{T_1} i\}\right) \\ &= \sum_{i=1}^{n} \sum_{j=i+1}^{n} \Big[\mathbf{1}\{i \prec_{T_1} j\} \left(\mathbf{1}\{j \prec_{T_2} i\} + \mathbf{1}\{i \perp_{T_2} j\}\right) + \mathbf{1}\{i \prec_{T_2} j\} \left(\mathbf{1}\{j \prec_{T_1} i\} + \mathbf{1}\{i \perp_{T_1} j\}\right) \\ &\quad + \mathbf{1}\{j \prec_{T_1} i\} \left(\mathbf{1}\{i \prec_{T_2} j\} + \mathbf{1}\{i \perp_{T_2} j\}\right) + \mathbf{1}\{j \prec_{T_2} i\} \left(\mathbf{1}\{i \prec_{T_1} j\} + \mathbf{1}\{i \perp_{T_1} j\}\right) \Big] \\ &= \sum_{i=1}^{n} \sum_{j=i+1}^{n} d^{i,j}(T_1, T_2). \end{aligned}$$

□

(Main Text) Lemma 3.4. *The AD distance $d(S, \mathcal{T})$ between a tree $S$ and trees $\mathcal{T}$ on the same vertex set $[n]$ equals*

$$d(S, \mathcal{T}) = \sum_{i=1}^{n} \sum_{j=i+1}^{n} d^{i,j}(S, \mathcal{T}) \tag{10}$$

*where $d^{i,j}(S, \mathcal{T})$ is the distance contributed by the unordered pair $\{i, j\}$ of distinct vertices defined as*

$$d^{i,j}(S, \mathcal{T}) = \mathbf{1}\{i \prec_S j\} \left(2a_{j,i} + b_{i,j}\right) + \mathbf{1}\{j \prec_S i\} \left(2a_{i,j} + b_{i,j}\right) + \mathbf{1}\{i \perp_S j\} \left(a_{i,j} + a_{j,i}\right). \tag{11}$$

Proof. We apply Lemma 3.1 and obtain

$$
\begin{aligned}
d(S,\mathcal{T}) &= \sum_{T\in\mathcal{T}} d(S,T) = \sum_{T\in\mathcal{T}}\sum_{i=1}^{n}\sum_{j=i+1}^{n} d^{i,j}(S,T)\\
&= \sum_{T\in\mathcal{T}}\sum_{i=1}^{n}\sum_{j=i+1}^{n} \big[\mathbf{1}\{i \prec_S j\}\,(2\cdot\mathbf{1}\{j\prec_T i\} + \mathbf{1}\{i \perp_T j\})\\
&\qquad + \mathbf{1}\{j \prec_S i\}\,(2\cdot\mathbf{1}\{i\prec_T j\} + \mathbf{1}\{i \perp_T j\})\\
&\qquad + \mathbf{1}\{i \perp_S j\}\,(\mathbf{1}\{i\prec_T j\} + \mathbf{1}\{j \prec_T i\})\,\big]\\
&= \sum_{i=1}^{n}\sum_{j=i+1}^{n}\left[\mathbf{1}\{i\prec_S j\}\left(2\sum_{T\in\mathcal{T}}\mathbf{1}\{j\prec_T i\} + \sum_{T\in\mathcal{T}}\mathbf{1}\{i\perp_T j\}\right)\right.\\
&\qquad + \mathbf{1}\{j\prec_S i\}\left(2\sum_{T\in\mathcal{T}}\mathbf{1}\{i\prec_T j\} + \sum_{T\in\mathcal{T}}\mathbf{1}\{i\perp_T j\}\right)\\
&\qquad \left. + \mathbf{1}\{i\perp_S j\}\left(\sum_{T\in\mathcal{T}}\mathbf{1}\{i\prec_T j\} + \sum_{T\in\mathcal{T}}\mathbf{1}\{j\prec_T i\}\right)\right] = \sum_{i=1}^{n}\sum_{j=i+1}^{n} d^{i,j}(S,\mathcal{T}).
\end{aligned}
$$

□

(Main Text) Lemma 4.2. *For any* $i, j \in V(\mathcal{T})$*, the entry* $a_{i,j}$ *of the ancestor-descendant matrix* $A_{\mathcal{T}}$ *equals:*

$$
a_{i,j} = \begin{cases}
2n^3 - 2n^2 + 4n - 3, & \text{if } 0 = i < j \le 2n,\\
n^3 - n^2 + 2n - 2, & \text{if } 0 < i < j \le n, (i,j) \in E(G),\\
n^3 - 2n^2 + 2n - 3, & \text{if } 0 < i < j \le n, (i,j) \notin E(G),\\
n^3 - n^2 + 2n - 1, & \text{if } 0 < i \le n < j \le 2n,\\
2n^3 - 2n^2 + 4n - 3, & \text{if } n \le i < j \le 2n,\\
0, & \text{if } 0 \le j < i \le 2n.
\end{cases} \tag{12}
$$

Proof. We prove the lemma by examining each of the six cases separately. For the first case, consider a pair $(i, j)$ such that $0 = i < j \le 2n$. Recall that $i = 0$ is the root vertex of all trees in $\mathcal{T}$. Thus, it holds that $i \prec_T j$ for any $T \in \mathcal{T}$. Therefore, $a_{i,j} = |\mathcal{T}| = 2n^3 - 2n^2 + 4n - 3$. For the second case, consider a pair $(i, j)$ such that $0 < i < j \le n$, $(i, j) \in E(G)$. Then, $i \prec_T j$ for all trees $T$ in $\mathcal{T}_0$ and $\mathcal{T}_i$. However, $i \not\prec_T j$ for any tree $T$ in the remaining multi-sets different from $\mathcal{T}_0$ and $\mathcal{T}_i$. Therefore, $a_{i,j} = |\mathcal{T}_0| + |\mathcal{T}_i| = (n^3 - 2n^2 + 2n - 3) + (n^2 + 1) = n^3 - n^2 + 2n - 2$. For the third case, consider a pair $(i, j)$ such that $0 < i < j \le n(i, j) \notin E(G)$. Then, $i \prec_T j$ for all trees $T$ in $\mathcal{T}_0$. However, $i \not\prec_T j$ for any tree $T$ in the remaining multi-sets different from $\mathcal{T}_0$ . Therefore, $a_{i,j} = |\mathcal{T}_0| = n^3 - 2n^2 + 2n - 3$. For the fourth case, consider a pair $(i, j)$ such that $0 < i \le n < j \le 2n$. Then, $i \prec_T j$ for all trees $T$ in the multi-sets $\mathcal{T}_0, \mathcal{T}_i$ and $\mathcal{T}_i'$. However $i \not\prec_T j$ for any tree $T$ in the remaining multi-sets. Therefore, $a_{i,j} = |\mathcal{T}_0| + |\mathcal{T}_i| + |\mathcal{T}_i'| = (n^3 - 2n^2 + 2n - 3) + (n^2 + 1) + 1 = n^3 - n^2 + 2n - 1$. For the fifth case, consider a pair $(i, j)$ such that $n < i < j \le 2n$. By construction, the chain $n + 1, \ldots, 2n$ is kept intact in every tree. Thus, $i \prec_T j$ for any tree $T \in \mathcal{T}$. Therefore, $a_{i,j} = |\mathcal{T}| = 2n^3 - 2n^2 + 4n - 3$. Finally, consider $(i, j)$ such that $0 \le j < i \le 2n$. For any $T \in \mathcal{T}$ and each edge $(i, j) \in E(T)$ it holds that $i < j$. Therefore, it holds that $i \not\prec_T j$ and thus $a_{i,j} = 0$ if $i > j$. □

(Main Text) Lemma 4.8. *For any pair* $(i, j)$ *of vertices such that* $0 \le i < j \le 2n$*, either* $i \prec_{S^*} j$ *or* $i \perp_{S^*} j$*.*

Proof. To prove this lemma, consider a tree $S$ such that $v <_S u$ for some $0 \leq u < v < 2n$. By Eq. (3) in Lemma 3.4, $d^{u,v}(S, \mathcal{T}) = 2a_{u,v} + b_{u,v}$. We distinguish three cases regarding the occurrence of $u$ and $v$, and show for each case that the resulting distance $d(S, \mathcal{T})$ will exceed the upper bound established in Lemma 4.7. First, consider $0 < u < v \leq n$. Then, $d^{u,v}_{\min}(\mathcal{T}) = a_{u,v}$ by Eq. (6), yielding

$$\begin{aligned} d(S,\mathcal{T}) = \sum_{i=0}^{2n} \sum_{j=i+1}^{2n} d^{i,j}(S,\mathcal{T}) &= d^{u,v}(S,\mathcal{T}) + \sum_{0<i<j\leq 2n,(i,j)\neq(u,v)} d^{i,j}(S,\mathcal{T}) \\ &\geq 2a_{u,v} + b_{u,v} + \sum_{0<i<j\leq 2n,(i,j)\neq(u,v)} d^{i,j}_{\min}(\mathcal{T}) \\ &= L + a_{u,v} + b_{u,v} = L + 2n^3 - 2n^2 + 4n - 3. \end{aligned}$$

Since $2n^3 - 2n^2 + 4n - 3 > n^2 - n$, Lemma 4.7, which states that $d(S^*, \mathcal{T}) \leq L + n^2 - n$, implies $S$ is not an optimal consensus tree.

Second, consider $0 < u \leq n < v < 2n$. Then, $d^{u,v}_{\min}(\mathcal{T}) = b_{u,v}$ by Eq. (6), yielding

$$\begin{aligned} d(S,\mathcal{T}) = \sum_{i=0}^{2n} \sum_{j=i+1}^{2n} d^{i,j}(S,\mathcal{T}) &= d^{u,v}(S,\mathcal{T}) + \sum_{0<i<j\leq 2n,(i,j)\neq(u,v)} d^{i,j}(S,\mathcal{T}) \\ &\geq 2a_{u,v} + b_{u,v} + \sum_{0<i<j\leq n,(i,j)\neq(u,v)} d^{i,j}_{\min}(\mathcal{T}) \\ &= L + 2a_{u,v} = L + 2n^3 - 2n^2 + 4n - 4 \end{aligned}$$

Since $2n^3 - 2n^2 + 4n - 4 > n^2 - n$, Lemma 4.7, which states that $d(S^*, \mathcal{T}) \leq L + n^2 - n$, implies $S$ is not an optimal consensus tree.

Third, consider $u = 0$ or $n < u < v \leq 2n$. Then $d^{u,v}_{\min}(\mathcal{T}) = b_{u,v} = 0$ by Eq. (6), yielding

$$\begin{aligned} d(S,\mathcal{T}) = \sum_{i=0}^{2n} \sum_{j=i+1}^{2n} d^{i,j}(S,\mathcal{T}) &= d^{u,v}(S,\mathcal{T}) + \sum_{0<i<j\leq 2n,(i,j)\neq(u,v)} d^{i,j}(S,\mathcal{T}) \\ &\geq 2a_{u,v} + b_{u,v} + \sum_{0<i<j\leq n,(i,j)\neq(u,v)} d^{i,j}_{\min}(\mathcal{T}) \\ &= L + 4n^3 - 4n^2 + 8n - 6 \end{aligned}$$

Since $4n^3 - 4n^2 + 8n - 6 > n^2 - n$, Lemma 4.7, which states that $d(S^*, \mathcal{T}) \leq L + n^2 - n$, implies $S$ is not an optimal consensus tree. □
